# Immature dendritic cells promote high-avidity tuning of vaccine T cell response

**DOI:** 10.1101/2020.07.16.204966

**Authors:** Adarsh Kumbhari, Colt A. Egelston, Peter P. Lee, Peter S. Kim

## Abstract

Therapeutic vaccines can elicit tumor-specific cytotoxic T lymphocytes (CTLs), but durable reductions in tumor burden require vaccines that stimulate high-avidity CTLs. Recent advances in immunotherapy responses have led to renewed interest in vaccine approaches, including dendritic cell vaccine strategies. However, dendritic cell requirements for vaccines that generate potent anti-tumor T-cell responses are unclear. Here we use mathematical modeling to show that counterintuitively, increasing levels of immature dendritic cells may lead to selective expansion of high-avidity CTLs. This finding contrasts with traditional dendritic cell vaccine approaches that have sought to harness ex vivo generated mature dendritic cells. We show that the injection of vaccine antigens in the context of increased numbers of immature dendritic cells results in a decreased overall peptide:MHC complex load that favors high-avidity CTL activation and expansion. Overall, our results provide a firm basis for further development of this approach, both alone and in combination with other immunotherapies such as checkpoint blockade.

## INTRODUCTION

In principle, the immune system can eliminate cancer cells by the activation and expansion of cancer-specific cytotoxic T lymphocytes (CTLs). Immune checkpoint blockade (ICB) immunotherapies, which release T cells from various negative regulatory pathways, have demonstrated impressive clinical successes and have become standard-of-care for many malignancies (1). However, the response to ICB seems to require the pre-existence of anti-tumor T cells (2). Vaccine approaches to generate tumor-specific T cells offer a potential solution towards generating a sufficient anti-tumor T cell response. Dendritic cell (DC) vaccines in particular, offer a means to activate and expand tumor-specific T cells (3). Here we discuss the impact of dendritic cell maturation status on vaccine design strategies.

CTLs detect cancer cells by T cell receptor (TCR) recognition of peptides displayed by a major histocompatibility complex (pMHC) on the surface of target cancer cells. Each TCR-pMHC interaction occurs at a particular strength – *affinity* – with multiple TCR-pMHC interactions occurring for each CTL-target cell interaction. While affinity is a measure of individual TCR-pMHC bonds, *avidity* is an overall measure of the strength of the TCR-pMHC interaction and more importantly determines the likelihood of successful lysis (4).

Therapeutic peptide vaccines aim to capitalize on the cancer-killing ability of CTLs. Initial results of peptide-based vaccines showed the ability to elicit significant numbers of antigen-specific CTLs, but often lacked measurable clinical successes (5-7). Recent progress in vaccine construction and combinatorial strategies with other immunotherapy agents have shown renewed promise for therapeutic peptide vaccines (3). Our work suggests that the dose and modality of peptide vaccines are key considerations for the design of future clinical interventions.

Early studies of cancer-specific CTLs showed that high-avidity TCRs are necessary to effectively lyse cancer cells that express native antigens at low levels (8). Preferentially selecting for high-avidity CTLs, however, is difficult. Regarding vaccines targeting cancer-associated antigens (CAA), thymic education of CTLs may likely have removed high-avidity T cells from the T-cell repertoire via negative selection (9). As a result, primarily low-avidity CTLs are left to respond to CAA-targeting vaccines. Beyond CAA, recent therapeutic vaccine efforts have focused on targeting somatic mutation derived neo-antigens (10, 11). As yet, neo-antigen vaccines have largely focused on the strength of peptide binding to MHC but have not yet explored the impact of dosage on T-cell repertoire response to the vaccine (12). For both CAA and neo-antigen targeting vaccines, standard dosages typically involve high antigen loads that may non-discriminately favor the expansion of both high and low avidity CTLs. However, lowering the dosage of peptides for vaccination yields sub-therapeutically relevant levels of CTL (13). Together, this highlights the need for further understanding of antigen dosage and context for efficacious vaccine design.

We previously showed that therapeutic vaccine designs were sensitive to dendritic cell-associated parameters (14). Given that DCs, which present antigen on their cell surface along with co-stimulatory molecules, facilitate CTL activation, we hypothesized that modulation of dendritic cell and peptide dosing could enhance an anti-cancer immune response. We show that by increasing the number of immature DCs, the average DC antigen load is lowered, which in turn selects for the expansion of high-avidity CTLs. This observation suggests traditional DC vaccine approaches that utilize ex vivo matured DCs may need to be reconsidered (3, 15). Our work suggests that combinatorial therapy with vaccine antigens and increased immature DCs, either by ex vivo generation or stimulated in vivo, may have efficacy. Thus, our findings suggest an approach that could improve already existing immune-based cancer therapies for increased and more durable clinical responses.

## MATERIAL AND METHODS

We previously developed a mathematical model to study how vaccine-induced avidity selection affects tumor clearance (14). This model was calibrated to ex vivo human data from Chung et al. (16) and then validated against data from (17, 18). Here, we extend this model to show that induction of immature DCs may improve current treatments by eliciting high-avidity CTLs. What follows is a brief description of our previously published model. We primarily use parameter estimates from the literature (see Table 1 and the references therein) and estimates generated from our prior analysis of ex vivo human data.

**Table 1.** Estimates that are characterized by human data are marked with a superscript H, while estimates based on murine data are marked with a superscript M. Additionally, estimates that are based on cell culture data are marked with a superscript V. Finally, the unit ‘k’ denotes 10^3^ cells.

| PARAMETER | DESCRIPTION | ESTIMATE | REFERENCE |
| --- | --- | --- | --- |
| $d_p$ | Peptide decay rate <sup>V</sup> | 6.16/day | (53) |
| $k_p$ | DC uptake rate <sup>HV</sup> | $3 \times 10^{-2}$ (k/μL)<sup>-1</sup>/day | (54) |
| $\delta_D$ | Immature DC decay rate <sup>HV</sup> | $5 \times 10^{-2}$/day | (55) |
| $s_D$ | Immature DC supply rate | $\delta_D M_{\text{total}}$ | Steady state |
| $M_{\text{total}}$ | Total DC population <sup>H</sup> | 23.4 k/μL | (56) |
| $d_D$ | Mature DC decay rate <sup>HV</sup> | 0.33/day | (55) |
| $\chi$ | Concentration of non-vaccine-associated proteins <sup>H</sup> | $6.72 \times 10^7$ ng/mL | (57) |
| $k_D$ | Mature DC presentation rate <sup>MV</sup> | $2.4 \times 10^5$ pMHCs/day | (58, 59) |
| $d_m$ | pMHC degradation rate <sup>MV</sup> | 2.9/day | (60) |
| $N$ | (Computational) maximum number of vaccine-associated pMHCs on a maturing DC | 700 | (14) |
| $J$ | Number of avidity levels | 20 | (14) |
| $s_T$ | Naive CTL supply rate | $d_N N(0)$ | Steady state |
| $s_H$ | Naive helper T cell supply rate | $d_{NH} N^H(0)$ | Steady state |
| $d_N$ | Naive CTL egress rate <sup>M</sup> | 1.2/day | (61) |
| $d_{NH}$ | Naive helper T cell egress rate <sup>M</sup> | 2.2/day | (61) |
| $N(0)$ | Initial naive CTL concentration <sup>M</sup> | $7.6 \times 10^{-3}$ k/μL | (62-65) |
| $N^H(0)$ | Initial naive helper T cell concentration <sup>M</sup> | 0.0571 k/μL | (62, 66) |
| $R_{LH}$ | Ratio of low-high avidity naive CTLs | 100 | Assumption |
| $k_{DC}$ | Naive CTL-DC interaction rate <sup>M</sup> | $0.4$ (k/μL)<sup>-1</sup>/day | (62) |
| $\tau_m$ | DC migration time <sup>M</sup> | 0.75 days | (62) |
| $V_{\text{tissue}}$ | Volume of tissue site | 1000 μL | (14) |
| $V_{LN}$ | Volume of lymph node <sup>M</sup> | 4.2 μL | (62) |
| $n_H$ | Number of helper T cell divisions <sup>HV</sup> | 10 | (67) |
| $\tau_a$ | T cell division time <sup>M</sup> | 1 day | (66, 68) |
| $d_H$ | Effector helper T cell decay rate <sup>H</sup> | 0.008/day | (69) |
| $n_T$ | Number of CTL divisions <sup>M</sup> | 15 | (68, 70-73) |
| $d_T$ | Effector CTL decay rate <sup>H</sup> | 0.009/day | (69) |
| $\varphi_0$ | Antigen saturation constant | $5 \times 10^3$ ng/mL | (14) |
| $r_1$ | Secretion rate of growth signal by CTLs | 0.1/day | (14) |
| $r_2$ | Secretion rate of growth signal by helper T cells | 1/day | (14) |
| $d_G$ | Growth factor decay rate <sup>H</sup> | 144.4/day | (74) |
| $k_G$ | CTL-growth factor interaction rate | $0.1$ (k/μL)<sup>-1</sup>/day | (14) |
| $k_R$ | Treg differentiation rate | 0.2/day | (14) |
| $d_R$ | Treg decay rate <sup>H</sup> | 0.083/day | (75, 76) |
| $\mu$ | CTL-Treg interaction rate <sup>H</sup> | $5$ (k/μL)<sup>-1</sup>/day | (14) |
| $K$ | (Computational) maximum number of cognate pMHCs expressed on cancer cell | 295 | (14) |
| $\gamma$ | Growth rate of melanomas <sup>H</sup> | 0.0185/day | (77-79) |
| $\kappa$ | Carrying capacity of melanomas <sup>M</sup> | 736 k/μL | (80) |
| $\alpha$ | pMHC regeneration rate <sup>MV</sup> | 8.4/day | (81, 82) |
| $k_T$ | Tumor-CTL interaction rate <sup>HV</sup> | $16.1$ (k/μL)<sup>-1</sup>/day | Estimate |
| $p_T$ | Probability of trogocytosis <sup>HV</sup> | 0.7 | Estimate |
| $\omega_1$ | Lysis likelihood for lowest avidity ($j=1$) CTL <sup>HV</sup> | 0.28 | Estimate |
| $\omega_J$ | Lysis likelihood for highest avidity ($j=J$) CTL <sup>HV</sup> | 0.96 | Estimate |
| $C_{\text{init}}$ | Initial cancer concentration | 0.05 k/μL | (14, 16) |

### Basic model

The model consists of three major components: the activation and maturation of dendritic cells (Equations 1 to 6); the activation and proliferation of T cells (Equations 7 to 12); and the lysis and trogocytosis-mediated MHC stripping of cancer cells by effector CTLs (Equations 17 to 19). Figure 1 depicts a schematic of these interactions.

**Figure 1.**
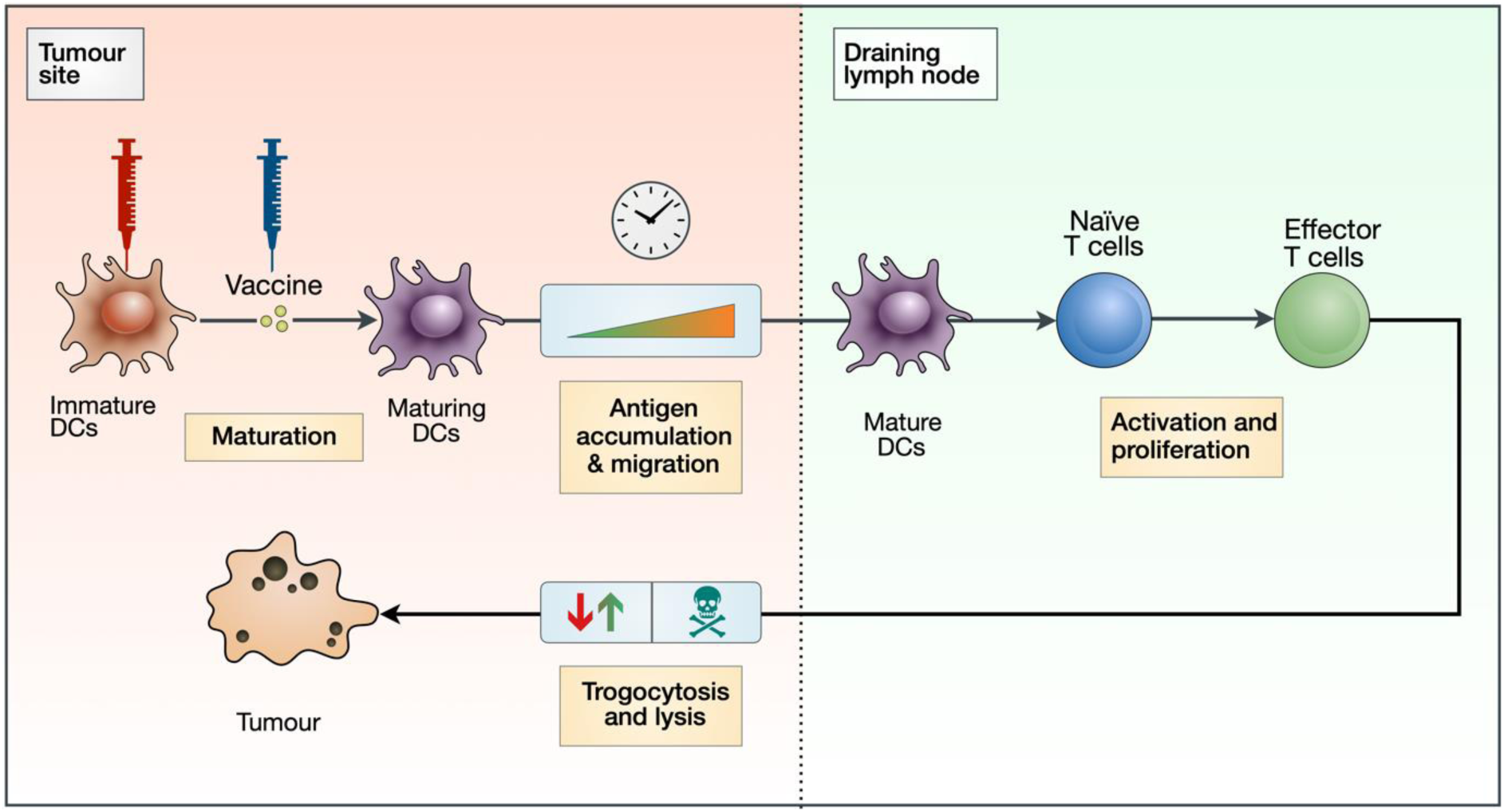
Block diagram depicting key aspects of our vaccination model. An injection of peptide vaccine activates immature DCs (which are also injected), prompting an accumulation of antigen by maturing DCs. These maturing DCs then migrate and activate naive T cells in the lymph node, which then proliferate into effector T cells. Effector T cells can both strip peptides off the surface of cancer cells via trogocytosis and kill cancer cells.

#### Dendritic Cells

To model the activation and maturation of dendritic cells (DCs) at the tissue site (the volume of which is *V*_tissue_), we consider several populations: *P*, the concentration of vaccine peptides; *I*, the concentration of immature DCs; and *M*_*j*_, the concentration of maturing DCs presenting *j* vaccine-associated pMHCs, where *j* can vary between zero and *N*. In modelling the interactions between these populations, we assume that immature DCs mature in the presence of peptide antigen and extracellular maturation signals. Dendritic cell maturation may occur in the presence of vaccine adjuvant, various danger signals, tissue derived immunogenic signals (19, 20). DC maturation signals may in turn affect T-cell priming and activation (17). As a simplifying assumption, we assume that the strategy to optimize DC maturation is successful. Next, we model the interactions between these populations with an ODE system:

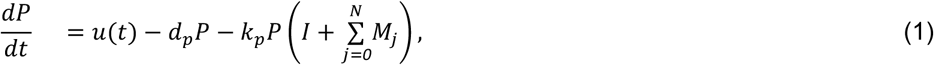

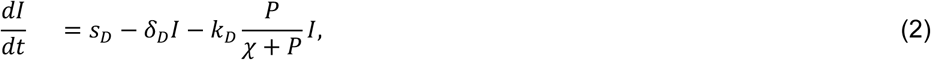

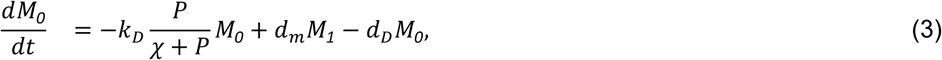

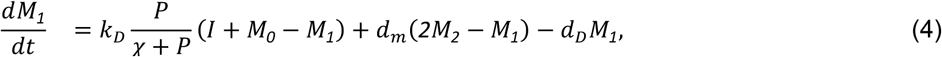

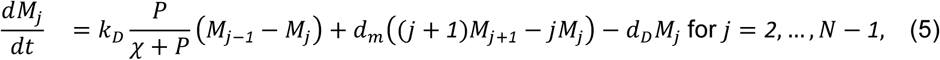

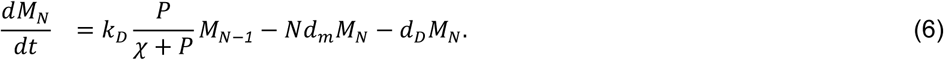

In Equation 1, vaccine peptides are injected at rate *u*(*t*), decay at rate *d*_*p*_, and as a simplifying assumption, taken up by both immature DCs and mature DCs at rate *k*_*p*_. In Equation 2, immature DCs are supplied at rate *s*_*D*_, and decay at rate *δ*_*D*_. Because of adjuvant, immature DCs are assumed to mature and acquire vaccine peptides at rate 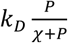. Here, *k*_*D*_ is the rate of peptide presentation, *χ* is the concentration of non-vaccine peptides, and 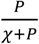 is the proportion of peptides presented that are vaccine specific.

In Equations 3 to 6, immature DCs initially enter the mature DC population presenting one vaccine peptide with subsequent peptides presented at rate 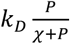 as described above. Additionally, surface peptides degrade at rate *d*_*m*_, which is proportional to the number of presented peptides, *j*. Finally, mature DCs decay at rate *d*_*D*_. Here, we assume that mature DCs decay faster than iDCs (21).

#### T Cells

To model the activation and proliferation of T cells both at the lymph node (the volume of which is *V*_LN_) and at the tissue site, we first model avidity as a spectrum that varies from *j*=1 to *j*=*J*, corresponding to the lowest and highest avidity states respectively. We then consider several populations: *N*_*j*_, the concentration of naive CTLs of avidity 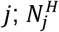, the concentration of naive helper T cells of avidity *j*; *T*_*j*_, the concentration of effector CTLs of avidity *j*; *H*_*j*_, the concentration of effector helper T cells of avidity *j*; *R*, the concentration of induced regulatory T cells; and *G*, the concentration of positive growth factors. The interactions between these populations are then modelled with an ODE system:

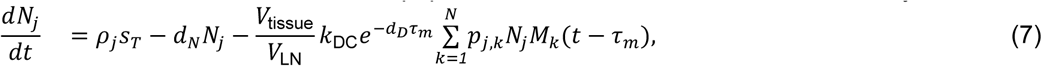

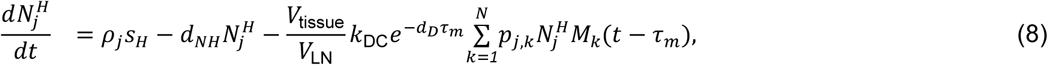

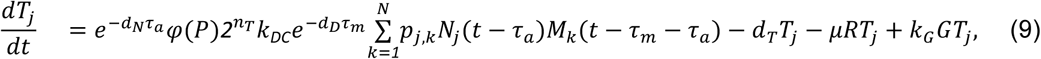

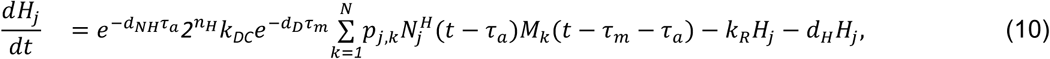

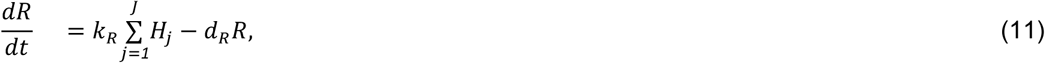

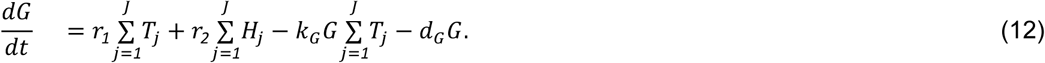

In Equation 7, naive CTLs in the lymph node of avidity *j* are supplied at rate *ρ*_*j*_*s*_*T*_, where *ρ*_*j*_ is the proportion supplied that have avidity *j*. These naive CTLs also exit the lymph node at rate *d*_*N*_. The rate at which naive CTLs are activated by mature DCs that have migrated into the lymph node is

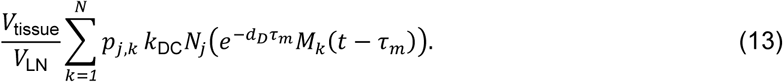

Migration is modelled with a fixed delay of *τ*_*m*_, with 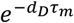 being the proportion that survives migration. The kinetic interaction rate between naive CTLs of avidity *j* and mature DCs presenting *k* vaccine-peptides is *k*_DC_ with *p*_*j,k*_ being the probability of an interaction leading to successful activation. Finally, the leading term 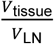 accounts for the volume change between the tissue site and the lymph node. In Equation 8, which is similar to Equation 7, naive helper T cells of avidity *j* are supplied at rate *ρ*_*j*_*s*_*H*_, decay at rate *d*_*NH*_, and are activated at the net rate of

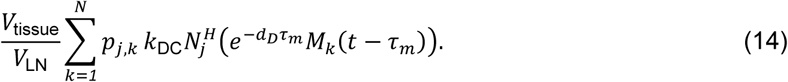

Equations 9 to 12 describe interactions within the tissue site. In Equation 9, naive CTLs undergo *n*_*T*_ divisions. The division program is modelled with a fixed delay of *τ*_*a*_, with 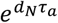 being the proportion that survives the division program, which equates to a net supply rate of

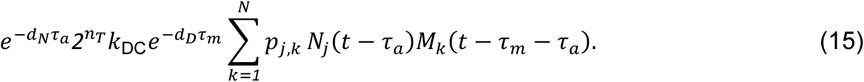

To account for T-cell hyporesponsiveness, we multiply Equation 15 by 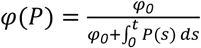. This ensures that antigen accumulation results in diminished effector CTL expansion. We also assume effector CTLs: decay at rate *d*_*T*_; expand due to interactions with positive growth factors at rate *k*_*G*_; and are suppressed by interactions with induced regulatory T cells at rate *μ*.

In Equation 10, naive helper T cells undergo *n*_*H*_ divisions. Following a similar argument to that in Equation 9, the net supply rate of effector helper T cells is

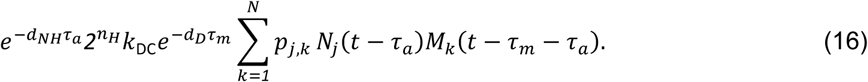

These effector helper T cells decay at rate *d*_*H*_ and differentiate into induced regulatory T cells at rate *k*_*R*_.

In Equation 11, regulatory T cells enter the system as differentiated effector helper T cells and decay at rate *d*_*R*_. Finally, in Equation 12, effector CTLs and helper T cells secrete growth factors such as IL-2 at rates *r*_*1*_ and *r*_*2*_. These growth factors are assumed to decay at rate *d*_*G*_.

#### Cancer cells

To model the lysis of cancer cells and trogocytosis of cancer cell MHC by effector CTLs, we consider a population of cancer cells presenting *k* vaccine-associated peptides, *C*_*k*_, where *k* varies from zero to *k*. The interactions between these cancer cells and effector CTLs are modelled with an ODE system:

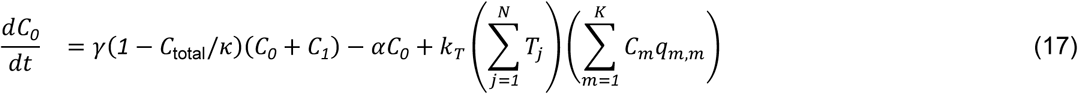

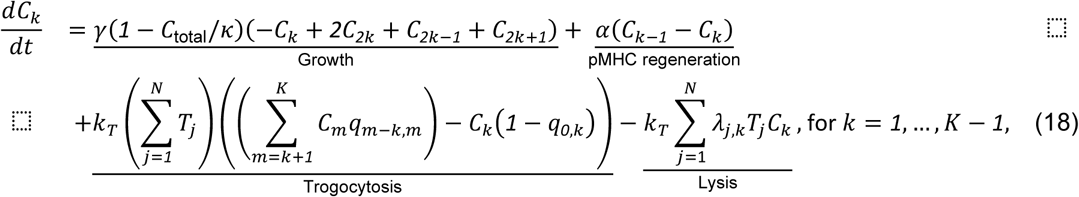

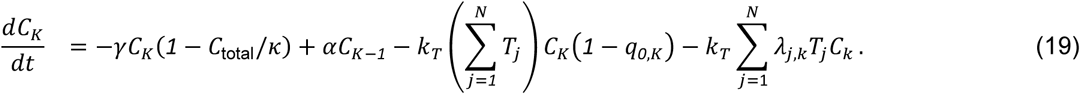

In Equations 17 to 19, the total cancer population, 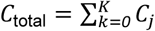, grows logistically at rate *γ* and with carrying capacity *κ*. As a simplifying assumption, we assume that the number of surface peptides is halved after mitosis, resulting in a net compartmental growth rate of

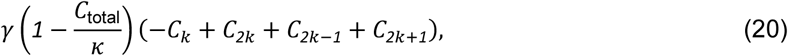

for the population of cancer cells presenting *k* peptides, *C*_*k*_. We also assume that surface peptides are regenerated at rate *α*. To model trogocytosis-mediated MHC stripping, we assume that CTLs and cancer cells presenting *k* peptides interact at rate *k*_*T*_ and additionally assume the number of peptides stripped during this interaction is binomially distributed with probability *p*_*T*_. For brevity we let 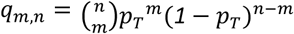 denote the probability that a CTL will trogocytose *m* MHC:peptides off a cancer cell presenting *n* surface peptides. This allows us to describe the trogocytosis rate as

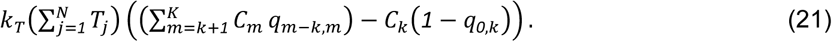

Finally, to model lysis, we let *λ*_*j,k*_ denote the lysis probability between a cancer cell presenting *k* peptides and an effector CTL of avidity *j*, and assume these interactions occur at rate *k*_*T*_. To model the lysis probability, we assume that the probability of lysis increases with cognate pMHCs but is also modulated by CTL avidity. This can be modelled by assuming a probability function of the form

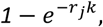

where *r*_*j*_ is an avidity-dependent rate parameter chosen so that the lysis probability at maximal levels of cognate pMHC expression, i.e., *λ*_*j,k*_ varies linearly from *ω*_*1*_ for the lowest avidity CTL to *ω*_*J*_ for the highest avidity CTL.

### Functional forms

#### Peptide vaccine injection rate

Here, we assume that the vaccine is injected systemically at a fixed dose, *u*_*0*_, and at a regular interval of ζ, which corresponds to the functional form

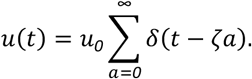

#### Activation probability

The probability of a mature DC presenting *k* vaccine-associated pMHCs activating a naive T cell of avidity *j, p*_*j,k*_, is modelled with a switch:

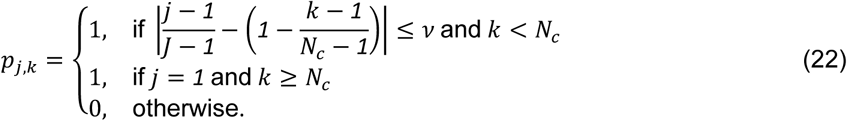

Here, 1/(*N*_*c*_ *−* 1) and 1/(*J −* 1) map *j* and *k* from their respective domains to [0,1]. The dimensionless parameter *ν* = 0.05 determines how sensitive our switching function is to pMHC expression. This characterization ensures that low pMHC levels on DCs stimulate high-avidity CTLs, and high pMHC levels on DCs stimulate both high- and low-avidity CTLs (18, 22-26). In contrast, low pMHC expression stimulates mostly low-avidity CTLs (9, 27-29). To reflect this, we assumed that beyond a critical number of pMHCs, *N*_*c*_, only low-avidity CTLs were stimulated. We set *N*_*c*_ = *N*/*2* = 350, implying that DCs must have *at least* a surface antigen density below 50% to stimulate high-avidity CTLs.

#### Initial Conditions

We assume that the vaccine is first administered at *t* = *0*, i.e., *P*(*0*) = *u*_*0*_, where *u*_*0*_ is the vaccine dose. To determine the initial DC populations, we assume that the system is at steady state when there is no vaccine, which implies *I*(*0*) = *M*_total_, and *M*_*j*_(*0*) = *0*, where *M*_total_ is the total DC population at steady-state conditions.

To model the scarcity of high-avidity naive T cells, we assume that their availability decreases exponentially. Specifically, we assume *N*_*j*_(*0*) = *ρ*_*j*_*N*(*0*) and 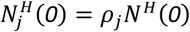, where *ρ*_*j*_ = *ae*^*−bj*^. Here, the model parameters *a* and *b* are chosen so that 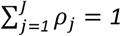 and *ρ*_*1*_ /*ρ*_*J*_, i.e., the ratio low-avidity to high-avidity T cells, equates to the model parameter *R*_LH_. In our simulations, we set *R*_LH_ to 100, which means that for one high-avidity T cell there are 100 low-avidity T cells.

For simplicity, we assume that initially there are zero vaccine-associated effector T cells, i.e., *T*_*j*_(*0*) = *0, H*_*j*_ (*0*) = *0*, and *R*(*0*) = *0*. As there are no vaccine-associated effector T cells present initially, we also set the concentration of growth factor to be *G*(*0*) = *0*.

Finally, we assume that the total cancer cell concentration is *C*_init_, with cognate pMHC being normally distributed with mean *μ* = 148 and variance *σ*^2^ = 49. Mathematically, if 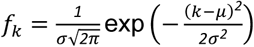, then 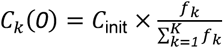.

### Sensitivity analysis

To understand how DC maturation status affects parameter sensitivity, we conduct sensitivity analysis on our modified model. We account for non-linear interactions between parameters by varying all parameters simultaneously using Latin hypercube sampling (*n*=250) over the ranges shown in Table 2, and measure sensitivity by calculating Spearman’s rank correlation coefficient (SRCC), ρ, for each parameter against the fold decrease. Table 2 shows SRCC ρ for each parameter.

**Table 2.** Spearman’s rank correlation coefficient between modified model parameters and fold decreases of simulations when varied simultaneously.

| PARAMETER | DESCRIPTION | RANGE | SRCC |
| --- | --- | --- | --- |
| $d_p$ | Peptide decay rate | $\pm 50\%$ | -0.0083 |
| $k_p$ | DC uptake rate | $\pm 50\%$ | -0.0462 |
| $\delta_D$ | Immature DC decay rate | $\pm 50\%$ | 0.0096 |
| $M_{\text{total}}$ | Total DC population | $\pm 50\%$ | 0.0508 |
| $d_D$ | Mature DC decay rate | $\pm 50\%$ | -0.0341 |
| $\chi$ | Concentration of non-vaccine-associated proteins | $\pm 50\%$ | 0.0596 |
| $k_D$ | Mature DC presentation rate | $\pm 50\%$ | -0.0357 |
| $d_m$ | pMHC degradation rate | $\pm 50\%$ | 0.0047 |
| $d_N$ | Naive CTL egress rate | $\pm 50\%$ | -0.0176 |
| $d_{NH}$ | Naive helper T cell egress rate | $\pm 50\%$ | -0.0096 |
| $N(0)$ | Initial naive CTL concentration | $\pm 50\%$ | 0.0884 |
| $N^H(0)$ | Initial naive helper T cell concentration | $\pm 50\%$ | 0.0801 |
| $R_{LH}$ | Ratio of low-high avidity naive CTLs | 10-500 | -0.1091 |
| $k_{DC}$ | Naive CTL-DC interaction rate | $\pm 50\%$ | 0.0545 |
| $\tau_m$ | DC migration time | $\pm 50\%$ | 0.0483 |
| $V_{\text{tissue}}$ | Volume of tissue site | $\pm 50\%$ | -0.0067 |
| $V_{\text{LN}}$ | Volume of lymph node | $\pm 50\%$ | 0.0529 |
| $n_H$ | Number of helper T cell divisions | 4-10 | -0.1187 |
| $d_H$ | Effector helper T cell decay rate | $\pm 50\%$ | 0.0527 |
| $\tau_a$ | T cell division time | $\pm 50\%$ | -0.0395 |
| $n_T$ | Number of CTL divisions | 10-20 | 0.4848 |
| $d_T$ | Effector CTL decay rate | $\pm 50\%$ | -0.0442 |
| $d_G$ | Growth factor decay rate | $\pm 50\%$ | 0.0461 |
| $d_R$ | Treg decay rate | $\pm 50\%$ | -0.0241 |
| $k_G$ | CTL-growth factor interaction rate | $\pm 50\%$ | 0.0454 |
| $k_R$ | Treg differentiation rate | $\pm 50\%$ | -0.0149 |
| $r_1$ | Secretion rate of growth signal by CTLs | $\pm 50\%$ | -0.0561 |
| $r_2$ | Secretion rate of growth signal by helper T cells | $\pm 50\%$ | 0.0028 |
| $\mu$ | CTL-Treg interaction rate | $\pm 50\%$ | -0.0662 |
| $\gamma$ | Growth rate of melanomas | 0.003- 0.087/d | -0.5958 |
| $\kappa$ | Carrying capacity of melanomas | 48.7-2360 k/$\mu\text{L}$ | 0.1457 |
| $\alpha$ | pMHC regeneration rate | $\pm 50\%$ | 0.1189 |
| $k_T$ | Tumor-CTL interaction rate | $\pm 50\%$ | 0.1761 |
| $p_T$ | Probability of trogocytosis | $\pm 50\%$ | -0.1905 |
| $\omega_1$ | Lysis likelihood for lowest avidity ($j=1$) CTL | $\pm 50\%$ | -0.0416 |
| $\omega_J$ | Lysis likelihood for highest avidity ($j=J$) CTL | $\pm 50\%$ | 0.3595 |

In our previous model, a sensitivity analysis identified antigen presentation by DCs as a key variable for the beneficial therapeutic value of vaccines. Here, we amend our model with the induction of immature DCs, resulting in supraphysiological levels of DCs. The resulting scale difference reduces the power of DC-associated parameters. Additionally, the model is now sensitive to the tumor growth rate, γ, suggesting that characteristics such as proliferative and apoptotic cell rates may affect the clinical response to the therapeutic vaccine.

## RESULTS

### Modified mathematical model

We previously found that the rate of antigen presentation by DCs determined the therapeutic value of an anti-tumor CTL response (14). Here, we hypothesize that inducing high levels of immature DCs would preferentially stimulate naive high-avidity CTLs by increasing the total concentration of mature DCs while lowering the average antigen density per DC. To test this proposed approach, we change Equation 2 in our original model (see Materials and Methods) to include a source term, *v*(*t*), which describes the elicitation of immature DCs, either by injection of ex vivo derived DCs or by recruitment of DC progenitors from the bone marrow via cytokine stimulation:

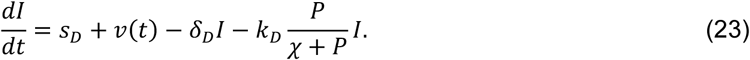

As a simplifying assumption, we assume that induced immature DCs (iDCs) are given at a fixed dose *v*_*0*_, and at dosing intervals of *ξ* hours *after* the injection of the peptide vaccine, which leads to the functional form:

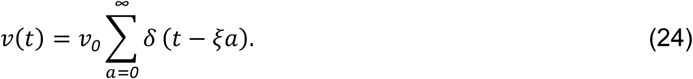

Figure 1 uses a block diagram to depict the key interactions of our model.

### Increased immature DC levels yields lower peptide:MHC levels and tumor cell reduction

In our example, we assume our tumor is a melanoma and assume that our vaccine either targets either neo-antigen peptides or classical antigens such as MART1. Initially, we simulate the DC context of the vaccine while leaving the peptide dosage fixed at the previously optimized value of 100 ng daily. Using this low peptide dosing, we effectively fix the pMHC levels on DCs to be low. To assess the robustness of our modified model, we next simulated iDC doses ranging from 10^3^ cells/μL to 10^12^ cells/μL, with dosing intervals that range from 0 to 24 hours after a peptide injection. A global sweep of iDC dosages within these ranges revealed an optimal iDC induction magnitude to be 5×10^5^ cells/μL of iDCs that induced a 98% decrease in tumor burden (Figure 2A). Importantly, the substantial reduction in tumor concentration we observed is neither dose dependent nor time dependent within our parameters, with a wide range of iDC concentrations and dosing intervals achieving a high degree of tumor reduction. Indeed, for iDC doses greater than 5×10^6^ cells/μL, the fold decrease in tumor concentration varies at most by 5% from the local optimum regardless of the dosing interval used. We thus find that the temporal robustness of this system centered around iDC induction and high-avidity T cell induction potentially allows for the possibility of introducing other combinatorial therapeutic strategies that may synergize with vaccine strategies, including checkpoint blockade and inducers of immunogenic cell death.

**Figure 2.**
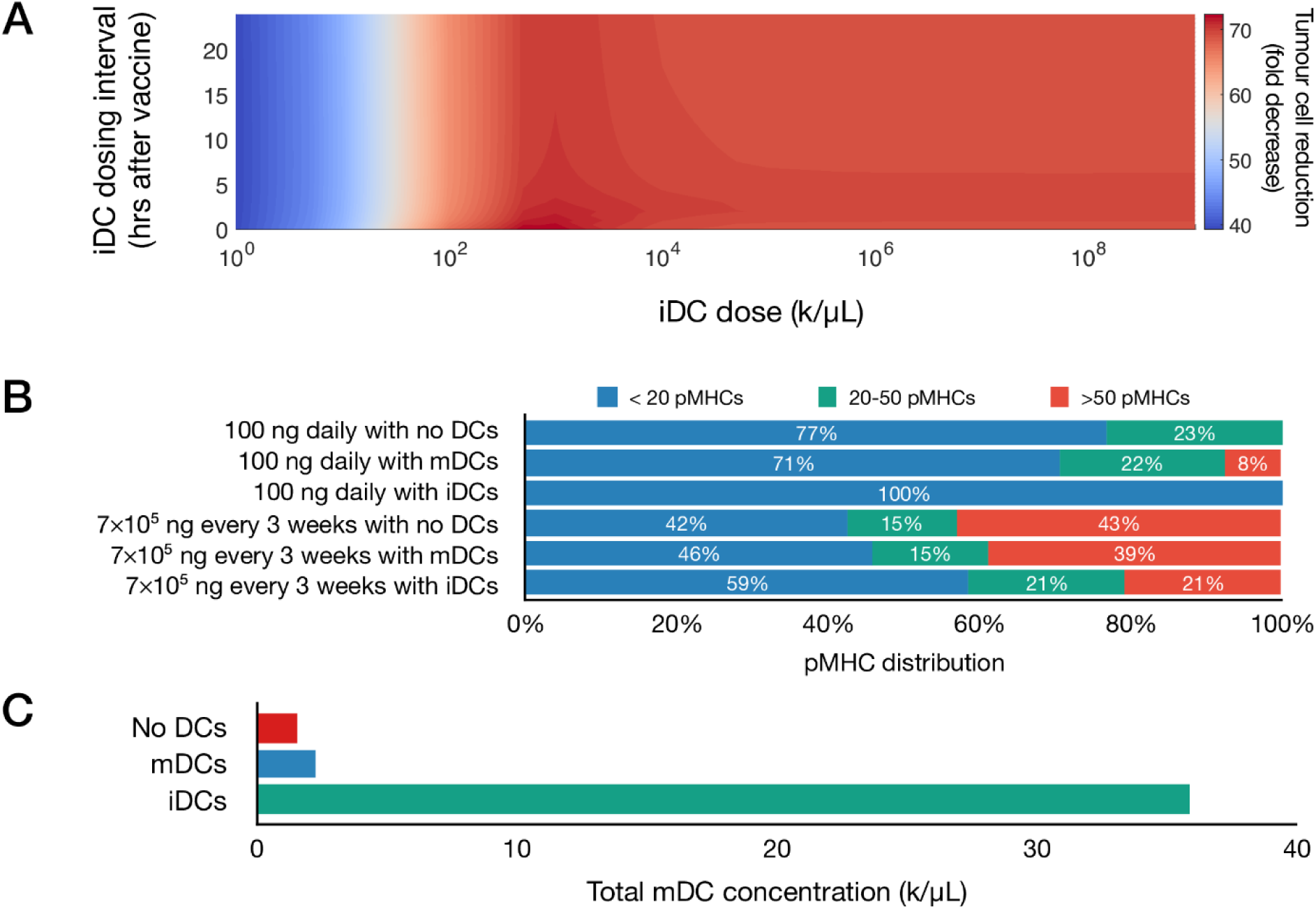
Induction of immature dendritic cells favors tumor reduction. **(A)** Heatmap depicts tumor cell reduction (fold change) for different iDC dosages when given with 100 ng of peptide daily. Here, the unit ‘k’ denotes 10^3^ cells. **(B)** Average distribution of antigen on mature DCs for various vaccine protocols. **(C)** Total concentration of mature DCs for various vaccine protocols using 7×10^5^ ng of peptide given every 3 weeks.

Our initial results demonstrated that increased iDC levels, rather than increased mDC levels, favor robust tumor clearing. We next set to determine if similar results could be recapitulated with clinically relevant vaccine dosages, rather than the 100 ng daily peptide dose identified by our model. We first compared pMHC levels in three therapeutic variations: peptide with either no DCs, induction of iDCs, or induction of mDCs with DC concentrations set to 6×10^3^ cells/μL, a dosing concentration previously used in a clinical setting and within optimal concentrations found in our global sweep above (30). We assume that within this population of ex vivo matured DCs (mDCs), pMHCs are normally distributed with mean *μ*=100 and variance *σ*^2^=25 (31). Additionally, we compare peptide dosing concentrations for both an ideal 100 ng daily and a clinically relevant 7×10^5^ ng every three weeks (18). Our model shows that at both peptide doses, induction of iDCs results in increased pMHC-low mature DCs as compared to no DC or mDC conditions (Figure 2B). This reduced antigen density in the context of the same peptide injection concentrations is due to the significantly increased numbers of mDCs generated by inducing iDCs (Figure 2C). These increased numbers are due to the longer half-life of iDCs as compared to mDCs, which are thought to rapidly decay upon maturation. As a result, the same peptide concentration dispensed over a larger number of DCs results in lower pMHC levels per DC.

### Immature DCs promote high-avidity T cells and tumor clearance in clinically relevant dosing schemes

Previously, we showed lower levels of pMHC competitively favor the expansion of high-avidity T cells rather than low-avidity T cells (14). As expected, we find that at both peptide dosing schemes induction of iDCs significantly favors the generation of high-avidity T cells compared to mDCs (Figure 3A). The optimal low dose of 100 ng daily of peptide significantly favors the development of high-avidity T cells, but even with the clinically relevant dosing of 7×10^5^ ng every three weeks, the induction of iDCs significantly shifts the balance of T cell composition to favor high-avidity T cells. This highlights that while traditional mDC or peptide-only vaccination strategies do increase T-cell induction, they do so at the expense of high-avidity T cells. In reflection of increased expansion of high-avidity T cells, our simulations further demonstrate that iDC induction results in improved cancer cell lysis (Figure 3B). Together, this suggests our iDC approach can be applied to current protocols to promote the expansion of high-avidity T cells and tumor clearance.

**Figure 3.**
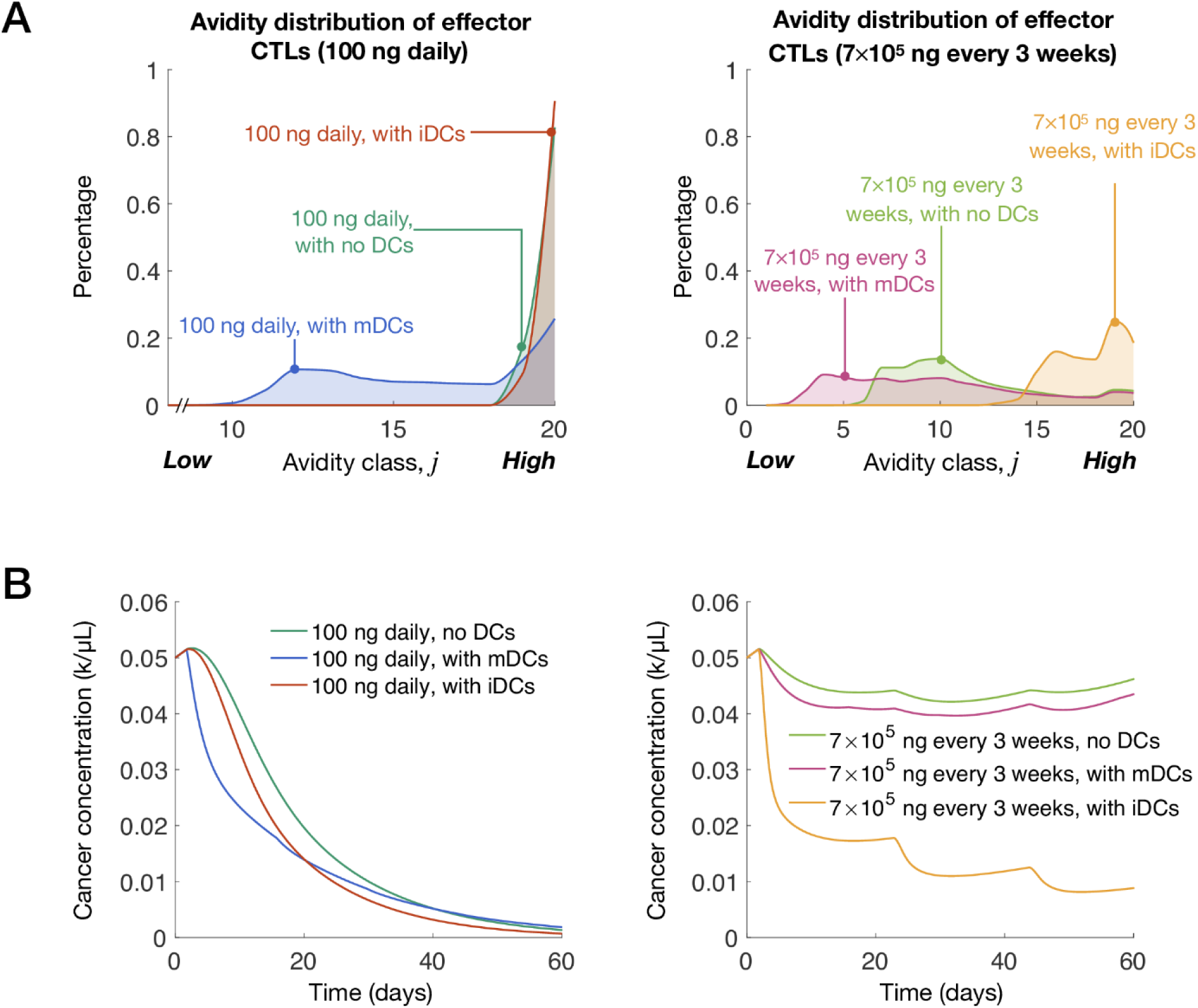
Induction of immature dendritic cells at clinically relevant vaccination doses yields significant tumor cell clearance. **(A)** Avidity distribution of effector T cells for various vaccine protocols. **(B)** Cancer concentrations over time for various vaccine protocols. The unit ‘k’ denotes 10^3^ cells.

## DISCUSSION

Cancer immunotherapy is now a routine means of successfully treating tumors of various types in the clinic. However, improved immunotherapies to benefit greater numbers of patients with increased durability are still needed. Despite its tremendous successes, ICB therapy only benefits less than the majority of patients treated (32-34) and presents significant risks for adverse side-effects (35-37). Therapeutic peptide vaccines can robustly induce a tumor-specific CTL response with limited side effects due to induction of an antigen-specific immune response rather than broad immune activation (18). Preferential development of high avidity anti-tumor CTLs enables enhanced tumor cell killing (8, 16). Previously, we showed that vaccine dosages could be optimized to preferentially elicit high-avidity CTLs, unlike standard dosages that elicit low-avidity CTLs (14). In that study, we showed that the efficacy of a dosage-optimized approach depended on DC-related parameters, which motivated us to explore how we could harness immature DCs to boost anti-tumor activity.

We hypothesized that increasing the magnitude of iDCs given with a dosage-optimized peptide vaccine may enhance CTL responses. It is important to stress that this approach is conceptually different from traditional DC vaccines in which ex vivo matured DCs are injected (3, 15). To assess this approach, we extended our previous model to account for a hypothetical induction of iDCs. We show that induction of iDCs, and not mDCs, can significantly reduce tumor burden, improving upon the performance of a peptide vaccine. A key assumption of our model is that iDCs will have a longer half-life and inducing iDCs will result in a larger overall pool of DCs as compared to the injection of mDCs, which are known to have a shorter half-life (21). Our simulations show that these effects are tied to the increased half-life of iDCs and therefore increased DC levels in general, which results in a lower average antigen density per DC. As such, induction of iDCs favors the preferential stimulation of high-avidity CTLs and tumor cell clearance.

Early cancer vaccines targeting over-expressed CAAs such as MART-1, MAGE, NYE-ESO-1, HER2, and MUC-1 demonstrated mediocre clinical results. Evidence suggests that the T cell repertoire capable of responding to these antigens are primarily composed of low-avidity T cells due to central tolerance of T cells specific for self-antigens (38). Recently, there has been renewed interest in cancer vaccines due to promising results for those targeting neoantigens (39-42). Additionally, encouraging preliminary clinical results have recently been observed in therapeutic approaches combining DC vaccines with checkpoint blockade (43). Our findings suggest that inducing increased iDC levels would benefit vaccines targeting either over-expressed CAAs or neoantigens, as the expansion of high-avidity CTLs would favor clinical responses in both scenarios.

Initial DC vaccines, such as Sipuleucel-T, were major milestones for immunotherapy-based treatments of cancer and demonstrated modest, but meaningful, clinical results (44). While DC vaccines have not achieved widespread therapeutic success, it is unclear if this is a result of targeting TAAs, the influence of previously unknown immunosuppression mechanisms in the tumor microenvironment, or difficult in manufacturing cell products (45). An exciting consequence of our findings is the concept that ex vivo maturation of autologous dendritic cells is an unnecessary, and possibly detrimental, step in vaccine design. Rather, in vivo induction of increased iDC levels, via strategies such as mobilization of bone marrow DC precursors, is an attractive possibility. Indeed, dosing with cytokines such as Fms-like tyrosine kinase 3 ligand (Flt3L) has demonstrated efficacy in increasing levels of circulating DCs (46-48). Finally, in support of our findings, increased circulating DC levels have been associated with increased survival in certain malignancies (49-51). Our model suggests that elevated levels of iDCs, rather than mDCs, favors a longer half-life of the circulating DC compartment and results in lower average pMHC levels that would then favor high-avidity T cell generation. Thus, induction of iDCs followed by peptide vaccination would favor tumor clearance. While we accept that maturation of iDCs is likely critical for tumor-specific T cell expansion, we suggest that in vivo maturation approaches may yield improved therapeutic results (52).

Our work addresses an important and less appreciated element of cancer vaccines – how vaccine design and administration can select for and enhance the proliferation of high-avidity CTLs. However, there remain many barriers to efficacy with a combination strategy that our model does not consider. For example, we do not account for potential intra-tumoral heterogeneity of antigen expression, factors influencing CTL trafficking to tumor sites, or a multitude of potential immune suppression mechanisms found within tumor microenvironments. Defining the minimum complexity of the immune system is challenging, and the model used in this study does not, nor does it aim to account for all known immune interactions.

The mathematical model presented here proposes that increasing the magnitude of immature DCs with an optimized peptide vaccine may improve tumor clearance. The model highlights the relative importance of antigen loads on dendritic cells, which facilitate the selective expansion of high-avidity CTLs. While pre-clinical experimental validation of our findings are necessary, our model suggests previously unappreciated aspects of vaccine design that may be necessary for the development of effective cancer treatments.

## AUTHOR CONTRIBUTIONS

All authors contributed to the conception and design of the study; AK performed the simulations and formal analysis; AK and CAE wrote the first draft of the manuscript.

